# Experience instructs the refinement of feature-selectivity in the mouse primary visual thalamus

**DOI:** 10.1101/2024.07.31.605460

**Authors:** Takuma Sonoda, Céleste-Élise Stephany, Kaleb Kelley, Di Kang, Rui Wu, Meghna R. Uzgare, Michela Fagiolini, Michael E. Greenberg, Chinfei Chen

**Affiliations:** Department of Neurology, F.M. Kirby Neurobiology Center, Boston Children’s Hospital, Boston, MA, United States; Department of Neurobiology, Harvard Medical School, Boston, MA, United States; Harvard-MIT Health Sciences and Technology Program, Harvard Medical School, Boston, MA, United States

**Keywords:** Synaptic plasticity, experience-dependent plasticity, thalamus, dLGN, retinogeniculate, circuit development

## Abstract

Neurons exhibit selectivity for specific features: a property essential for extracting and encoding relevant information in the environment. This feature-selectivity is thought to be modifiable by experience at the level of the cortex. Here, we demonstrate that selective exposure to a feature during development can instruct the population representation for that feature in the primary visual thalamus. This thalamic plasticity is not simply inherited from the cortex because it is still observed when recordings were performed in the absence of cortical feedback. Moreover, plasticity is blocked in mutant mice that exhibit deficits in retinogeniculate refinement, indicating that alterations in feature-selectivity are a direct result of changes in feedforward connectivity. These experience-dependent changes persist into older ages—highlighting the importance of this developmental period in shaping population coding in the thalamus. Our results show that salient environmental features are hard-wired into thalamic circuits during a discrete developmental window.

## Introduction

Sensory experience plays an important role in refining synaptic connections during development. A powerful model system for investigating the mechanisms of experience-dependent circuit refinement is the mouse retinogeniculate synapse. Once the axons of retinal ganglion cells (RGCs) form synapses with relay neurons of the dorsal lateral geniculate nucleus (dLGN), the synapses are pruned and strengthened by a process that initially depends on spontaneous activity^1–5^ and is associated with a sharpening of the geniculate receptive fields^6,7^. In mice, this initial phase of synaptic refinement and receptive field sharpening is followed by a second period of experience-dependent plasticity (postnatal days (P) 20-35) during which retinogeniculate connectivity is rewired^8,9^. While this rewiring requires visual experience, this plasticity does not result in a significant change in the receptive field sizes of dLGN neurons^7^, raising the question of why it occurs.

Recent studies in mice have shown that, even in the adult, a single dLGN neuron can be innervated by many RGCs^10–12^. Moreover, RGCs converging onto a single dLGN neuron can consist of multiple types that encode distinct visual features^13–16^. RGC types tile visual space independently of one another^17,18^, prompting the hypothesis that experience instructs the competitive refinement of RGC types. In this model, inputs from RGC types that respond to salient features in the environment are maintained and strengthened while inputs from RGC types that encode less prevalent features are functionally pruned. This refinement would be predicted to occur without any overall alterations in dLGN receptive field size. To test this hypothesis, we restricted the visual experience of animals to bias the activity of certain RGC types. If experience instructs the types of inputs that are selected and strengthened, we predicted that this manipulation would alter the tuning properties of downstream dLGN neurons to reflect features in the environment.

Here, we restricted the visual experience of animals to horizontal gratings moving upward during the experience-dependent phase of retinogeniculate refinement (P20-35). In response to selective exposure to this stimulus, we find the proportion of dLGN neurons that prefer the vertical axis of motion increases. These results show that experience instructs retinogeniculate refinement in a feature-specific manner and reveal a surprising degree of plasticity in the tuning properties of thalamic neurons.

## Results

### Selective Experience Rearing (SER) alters the tuning properties of dLGN neurons

We tested whether experience could alter feature-selectivity by restricting the visual experience of animals using a selective experience rearing (SER) protocol in which P20 mice were placed in a chamber surrounded by 4 monitors for ∼3 hours per day and kept in total darkness for the remainder of the day (Figure 1D). Visual stimuli during SER consisted either of horizontal square-wave gratings moving upward (vertical SER or vSER), or gratings moving alternately in 8 different directions (8 direction or 8dSER). The spatial frequency of the gratings was 0.04 cycles/degree measured from the center of the SER chamber and the temporal frequency was 2 cycles/second. Normally reared (NR) mice were kept on a 12-hour light cycle in standard housing conditions.

**Figure 1:**
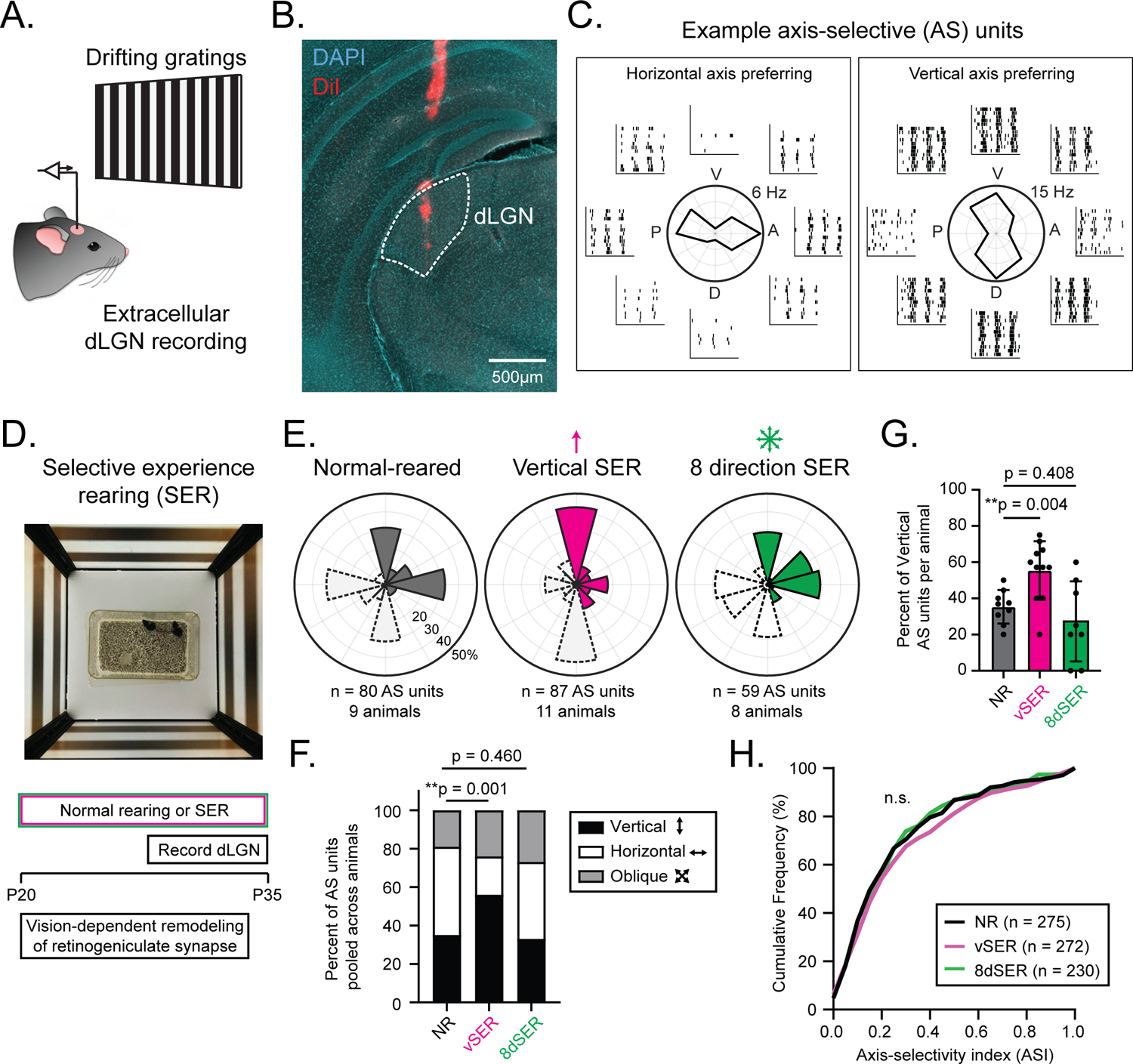
Selective experience rearing (SER) alters the tuning properties of dLGN neurons. **(A)** Schematic depicting *in vivo* dLGN recordings from anesthetized mice. Animals were presented with drifting sine-wave gratings moving in different directions to measure axis-selectivity (AS). **(B)** Example coronal brain section collected after *in vivo* recordings. The electrode tract trajectory was visualized by coating the recording electrode with DiI (red). This example shows the electrode passing through the dLGN (white dotted boundary). **(C)** Example recordings from two AS units in the dLGN. In the middle is a polar plot with average responses to drifting gratings moving in 8 different directions. Raster plots surrounding each polar plot represent spiking activity in response to gratings moving in different directions across 12 trials. A (anterior), V (ventral), P (posterior), D (dorsal). **(D)** Top: picture of the chamber used for SER. Mice in their home cage were placed in a chamber surrounded by 4 monitors for 3 hours per day. Bottom: timeline of SER experiments. Mice were subjected to either in normal rearing (NR) conditions or SER between P20-35. dLGN recordings were performed between P27-35. **(E)** Polar histograms showing the distribution of preferred axes of AS dLGN neurons in response to different rearing conditions. Preferred axes of AS units are plotted (as opposed to direction), with data points ranging from 0-180 degrees. Bins with dotted borders are mirror images intended to illustrate preferred axes of motion. **(F)** Proportion of units that prefer vertical (black), horizontal (white), or oblique (gray) motion. Units were pooled across animals. vSER significantly altered the distribution of preferred axes in AS dLGN neurons. **p < 0.01, Χ2 test. **(G)** Average percentage of AS dLGN units that prefer the vertical axis of motion measured across animals. Each data point represents a single animal. ** p < 0.01, Mann-Whitney test. **(H)** Cumulative distribution of ASI from all visually responsive units. SER did not result in any significant changes in ASI. n.s. (not significant), Kolmogorov-Smirnov test.

Following either NR or SER for 7-15 days, we recorded extracellularly from dLGN neurons in anesthetized mice and measured responses to drifting sine-wave gratings (Figure 1A-B). We focused on axis-selective (AS) neurons in the dLGN, which respond optimally to motion along a specific axis (Figure 1C) and constitute a significant proportion of dLGN neurons^19–23^. AS neurons were identified as having an axis-selectivity index (ASI) greater than 0.33 (see methods). We asked whether the distribution of preferred axes within the AS population changes depending on the nature of the visual experience. In NR mice, the distribution of preferred axes was evenly distributed between the horizontal and vertical axes (Figure 1E). In experimental animals that underwent vSER, the proportion of dLGN units that prefer the vertical axis of motion increased significantly when comparing the distribution of preferred axes of AS units pooled across animals (Χ^2^ test, p = 0.001) (Figure 1E-F). 8dSER did not result in a significant change in the distribution of preferred axes (Χ^2^ test, p = 0.460) (Figure 1E-F), indicating that changes we see in the vSER condition are not due to differences in light exposure (3 hours/day in SER vs. 12 hours/day in NR conditions).

To examine variability across animals, we measured the proportion of AS units that prefer vertical motion in each animal (Figure 1G). The percentage of units that prefer vertical motion was identical when comparing averages measured from AS units pooled from all animals with averages measured across animals (35% pooled vs. 35% averaged from 9 animals in NR and 56% pooled vs. 55% averaged from 11 animals in vSER) (Figure 1F-G). Importantly, significant changes in vSER animals were still observed when we compared the proportion of AS units preferring vertical motion across animals (Figure 1G). Additionally, we did not observe any alterations in the cumulative distribution of ASI in visually responsive neurons recorded from SER animals compared to NR (Figure 1H), suggesting that SER does not alter the proportion of AS dLGN neurons.

To determine whether dLGN receptive field sizes were impacted by SER, we measured the spatial receptive fields of dLGN neurons using spike-triggered averaging of responses to white noise movies^23,24^. vSER did not alter the receptive field sizes (Figure S1A-B) or the proportion of visually responsive dLGN neurons (Figure S3) when compared to NR mice. The locations of single-unit receptive fields also did not differ between NR and vSER animals, indicating that SER-dependent changes in axis preference were not due to differences in recording location (Figure S1C). Additionally, the distribution of ON-OFF responses recorded from visually responsive dLGN neurons to full-field flashes was not significantly different between vSER and NR animals (Figure S1D-E). Therefore, vSER specifically altered the preferred axes of AS dLGN neurons without disrupting spatial receptive field sizes or the distribution of ON-OFF responses of dLGN neurons. Taken together, these data show that visual experience shapes feature-selectivity in the developing dLGN.

### SER-dependent changes are not due to alterations in cortical feedback

Because the window of plasticity we observe in thalamus overlaps with the critical period in visual cortex^25–27^, we asked whether observed changes in dLGN receptive fields reflect underlying alterations in cortical feedback connections. We therefore sought to acutely silence visual cortex and measure the effect on dLGN receptive fields. Cortical silencing was achieved by expressing excitatory DREADDs in cortical inhibitory neurons^28^ with a local injection of AAV2/9-hDlx-GqDREADDs between P1-3 (Figure 2A). Later subcutaneous injections of clozapine-N-oxide (CNO) in P27-40 mice silenced the cortex within 10 minutes and lasted for at least 2 hours, as validated by multiunit recordings in response to drifting grating stimuli (Figure 2B-C). Expression of GqDREADDs was homogeneous across cortical layers (Figure S2A-B) and robust silencing was observed across the entire depth of the multielectrode array (Figure S2C). We also performed current source density analysis^24^ to confirm that infragranular layers that provide corticothalamic inputs were effectively silenced by activating Gq-DREADDs (Figure S2D).

**Figure 2:**
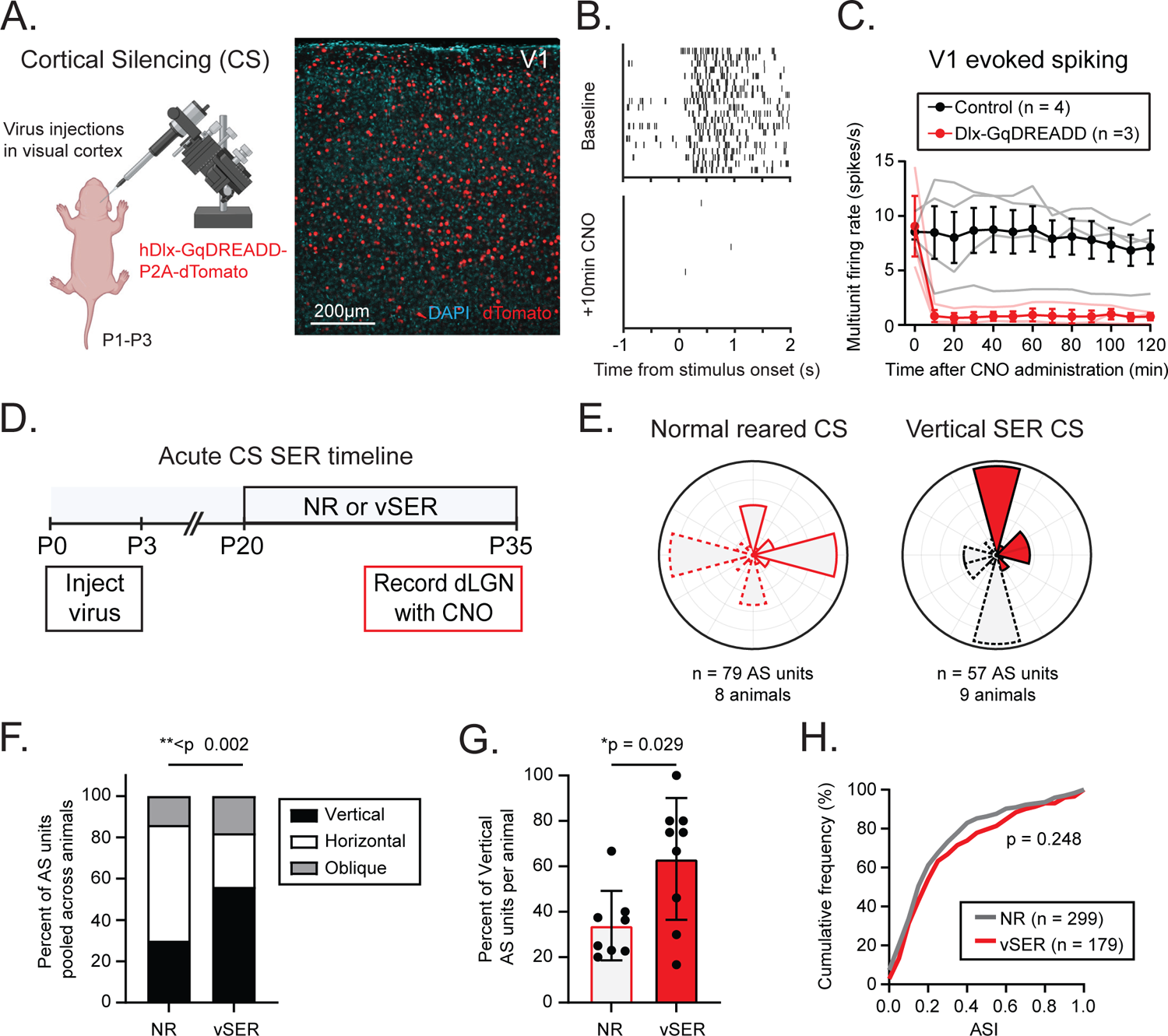
SER-plasticity is not due to alterations in cortical feedback connections. **(A)** Left: schematic depicting cortical injections of AAV9/hDlx-GqDREADD-P2A-dTomato in mouse pups between P1-3. Right: Example image from the primary visual cortex (V1) showing expression of Gq-DREADDs (red) driven by the hDlx promoter. **(B)** Raster plot from a V1 recording just prior to (top) and 10 minutes post CNO injection (bottom). Responses are multiunit activity recorded from a single channel in response to full-field drifting grating stimuli. **(C)** Grouped data showing average evoked activity over a 2-hour span after CNO injection. Control animals did not receive cortical injections of AAVs and were injected with CNO during cortical recordings. **(D)** Timeline of acute cortical silencing experiments. Cortex was silenced during dLGN recordings to block cortical feedback. **(E)** Polar histograms showing the distribution of preferred axes of AS dLGN neurons in experiments where cortex was acutely silenced for mice that were NR (left) or underwent vSER (right). **(F)** Proportion of units that prefer vertical (black), horizontal (white), or oblique (gray) motion pooled across animals. vSER mice exhibit significant increases in the number of AS units that prefer vertical motion in the absence of cortical feedback. p < 0.002, Χ^2^ test. **(G)** Average percentage of AS dLGN units that prefer the vertical axis of motion measured across animals. *p = 0.029, Mann-Whitney test. **(H)** Cumulative distribution of ASI from all visually responsive units. SER did not result in any significant changes in ASI. p = 0.248, Kolmogorov-Smirnov test.

We then made dLGN recordings from animals that received cortical injections of hDlx-GqDREADD virus (Figure 2D). CNO was administered subcutaneously at least 10 minutes prior to data collection to eliminate the cortical contributions to dLGN receptive field properties. Our results show that in the absence of cortical feedback, AS dLGN neurons still exhibit a significant increase in preference for vertical motion in vSER animals compared to NR animals (Figure 2D-G). Therefore, SER-dependent changes in dLGN receptive field properties are not due to changes in corticothalamic feedback.

### SER-dependent changes require proper retinal refinement

Another potential mechanism for SER-plasticity is that experience instructs the refinement of convergent RGC inputs, resulting in changes in dLGN neuron tuning. To test this possibility, we utilized mutant mice that carry 4 mutations in the methyl-CpG-binding 2 (MeCP2) gene (referred to as “MeCP2 Quadruple Knock-in” or “MeCP2 QKI” mice)^29^. These 4 mutations convert 4 important sites of activity-dependent phosphorylation on the MeCP2 protein to alanines. Using electrophysiological recordings in dLGN brain slices, we have previously shown that synaptic development is initially normal at P14 in MeCP2 QKI mice; however, between P14-30, MeCP2 QKI mice exhibit deficits in retinogeniculate refinement^29^ (Figure 3A). Therefore, MeCP2 QKI mice were used as a model to understand how SER-plasticity is altered in the absence of normal retinogeniculate refinement. If SER-dependent alterations are due to feature-specific alterations in RGC inputs, we predicted that alterations that occur in response to vSER would be disrupted in MeCP2 QKI mice. Indeed, AS dLGN neurons in MeCP2 QKI mice exposed to vSER do not exhibit any significant changes in the distribution of preferred axes (Figure 3B-D). These results show that normal retinogeniculate refinement is required for SER-dependent changes to occur, suggesting that SER ultimately exerts its influence on retinal connectivity in the dLGN.

**Figure 3:**
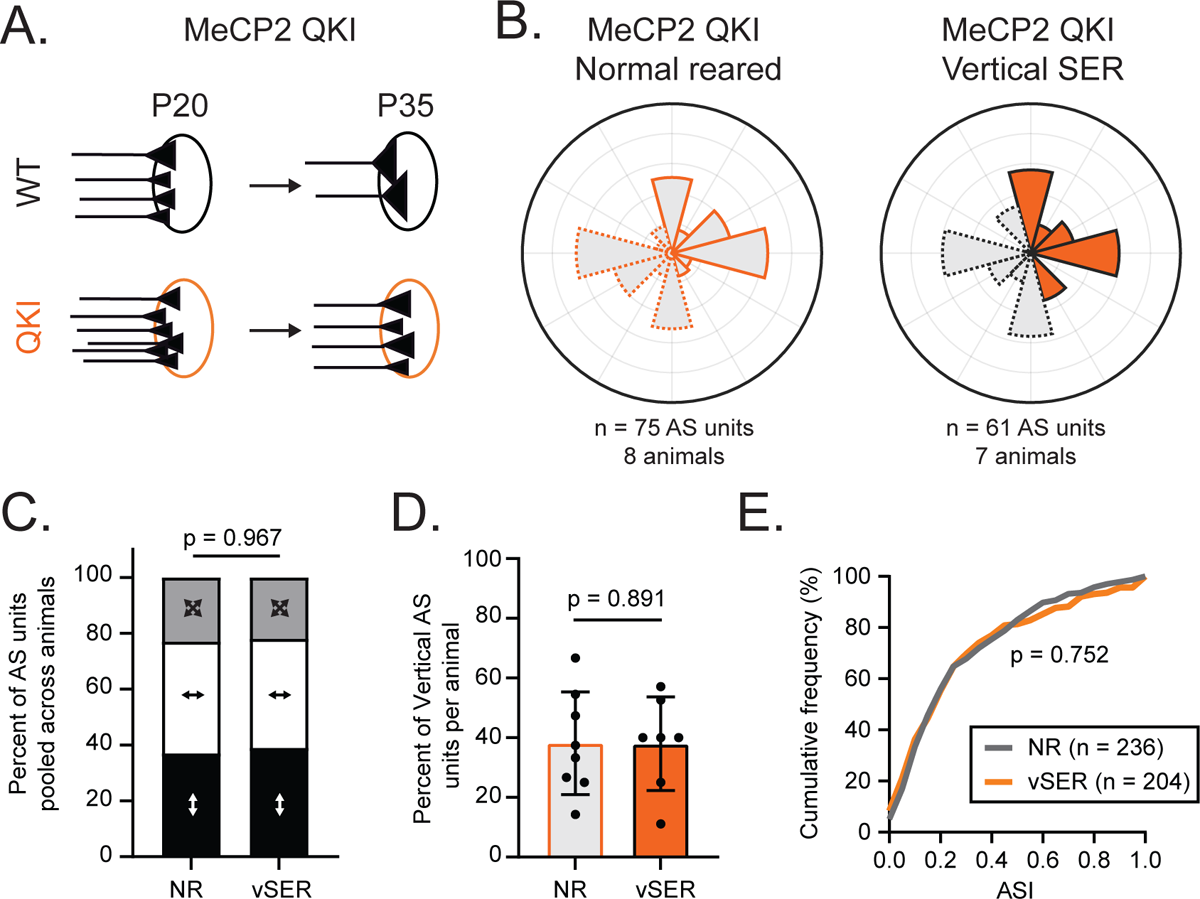
SER-plasticity requires proper retinogeniculate refinement. **(A)** Schematic depicting deficits in retinogeniculate refinement in MeCP2 QKI mice (Tzeng et al., 2023). **(B)** Polar histograms showing the distribution of preferred axes of AS dLGN neurons in MeCP2 QKI mice that were NR (left, gray) or underwent vSER (right, orange). **(C)** Proportion of units that prefer vertical (black), horizontal (white), or oblique (gray) motion pooled across animals. SER does not alter the distribution of preferred-axis in the dLGN of MeCP2 QKI mice. p = 0.967, Χ^2^ test. **(D)** Average percentage of AS dLGN units that prefer the vertical axis of motion measured across animals. P = 0.891, Mann-Whitney test. **(E)** Cumulative distribution of ASI from all visually responsive units. p = 0.752, Kolmogorov-Smirnov test.

### SER-dependent changes are maintained in adult mice

We next asked whether experience-dependent changes in dLGN receptive fields persist in adult mice. To test this, we returned mice to NR conditions after P20-35 SER and recorded from dLGN neurons between P60-75 (Figure 4A). Overall, the preferred axes of AS dLGN neurons maintained a bias towards vertical motion in vSER mice compared to NR mice after a 25-40 day “recovery” period when we compared units pooled across animals (Χ^2^ test, p = 0.005) (Figure 4B-C) and when we compared the proportion of vertical AS units across animals (Mann-Whitney test, p = 0.027) (Figure 4D). AS dLGN neurons in animals that underwent 8dSER did not exhibit any significant changes (Figure 4B-D). There was also no difference in the distribution of ASI in SER animals compared to NR (Figure 4E). These data indicate that stable representations of environmentally relevant features are formed in the dLGN during the experience-dependent phase of retinogeniculate refinement and retained into adulthood.

**Figure 4:**
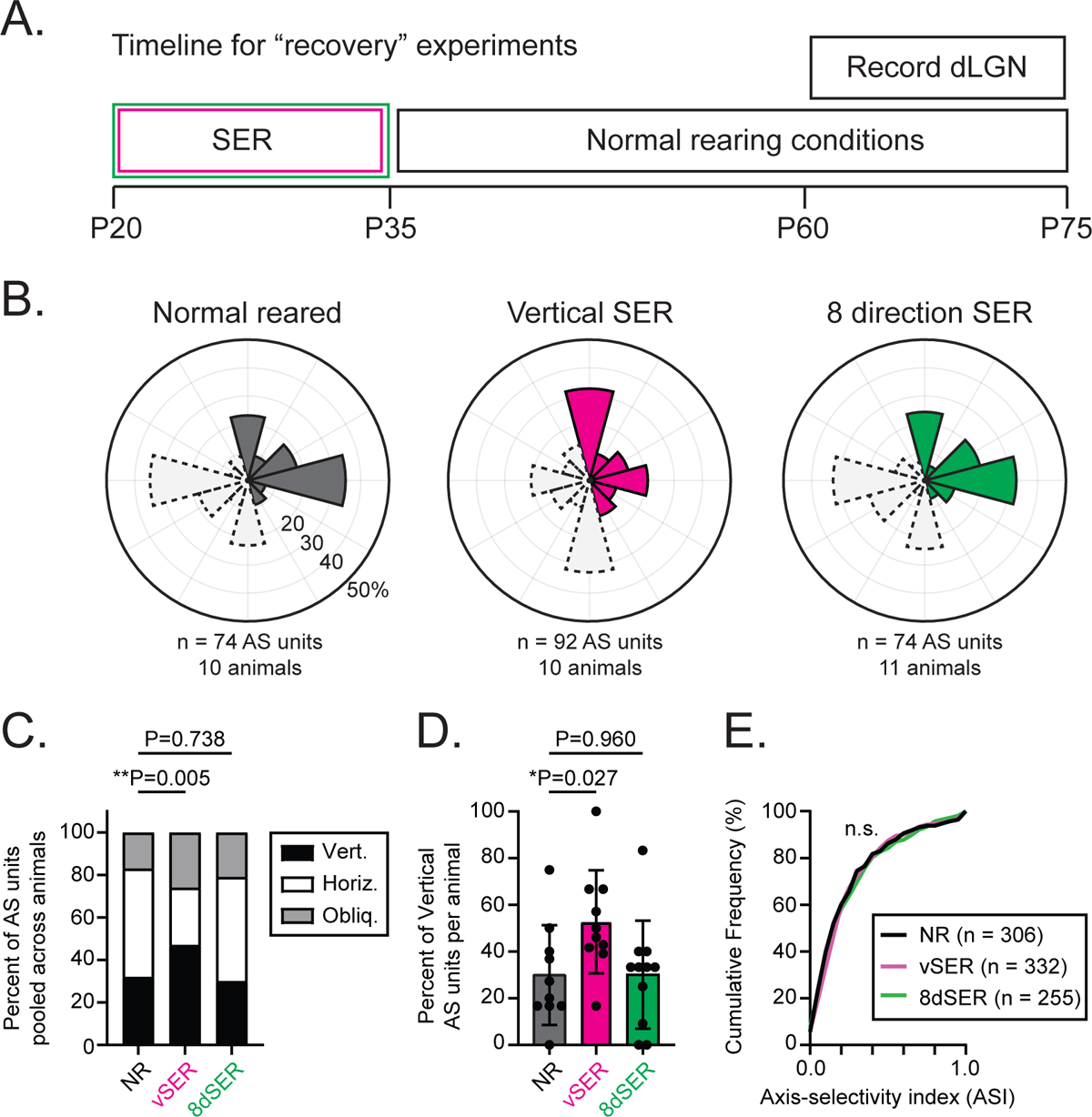
Changes acquired during SER are maintained in adult mice. **(A)** Timeline of recovery experiments in which animals were returned to normal rearing conditions between P35-75 after SER. dLGN recordings were performed between P60-75. **(B)** Polar histograms depicting the distribution of preferred axes of AS dLGN neurons in recovery experiments. Animals that underwent vSER maintain a vertical bias. **(C)** Proportion of units that prefer vertical (black), horizontal (white), or oblique (gray) motion. **p < 0.01, Χ^2^ test. **(D)** Average percentage of AS dLGN units that prefer the vertical axis of motion measured across animals. *p < 0.05, Mann-Whitney test. **(E)** Cumulative distribution of ASI from all responsive units. n.s. not significant, Kolmogorov-Smirnov test.

## Discussion

Our findings demonstrate that environmental features shape population coding in the mouse dLGN, reinforcing growing evidence for experience-dependent plasticity in the dLGN^8,30–34.^ This study is the first to show that features in an animal’s rearing environment—not just the absence or presence of visual input—can instruct features encoded by thalamic neurons.

Our results show that the encoding of axis of motion can be altered by visual experience in the dLGN during late postnatal development. This change in distribution cannot be explained simply by a decrease in the responsiveness of horizontal AS units because we observed no significant change in the proportion of visually responsive units in vSER animals (Figure S3).

Moreover, we observed no difference in the proportion of AS units in vSER animals (29% in NR vs 32% in vSER) in addition to a slight rightward shift in the cumulative distribution of ASI (Figure 1H). These results support the idea that experience instructs the axis preference of mouse dLGN neurons—unlike one model in cat visual cortex where experience maintains existing axis preference^35^. However, it is important to note that our results cannot rule out a scenario in which an increase in vertical AS units occurs simultaneous to an equal and opposite decrease in horizontal AS units.

Alterations we observe in the dLGN are unlikely inherited from retina. Population tuning of direction-selective and orientation-selective RGCs are unaffected by dark-rearing^36,37^ and direction preferences of motion-selective RGCs are unaltered by selective exposure to moving stripes^38,39^. While Zhang et al.^39^ did find that motion training, which included moving dots and gratings, improved the response reliability of on-off direction selective RGC responses to lower contrast stimuli, our study only utilized high-contrast (100%) gratings for both SER and dLGN recordings. Therefore, it was not possible to assess changes in response reliability at lower contrasts. In the future, it will be important to determine how experience-dependent changes in response reliability alter postsynaptic responses in dLGN neurons and how other features, such as contrast, can be fine-tuned at the level of the dLGN.

Analogous to long-term potentiation^40^, SER-plasticity may rely on distinct mechanisms for its triggering and expression. Our data suggest that changes in retinogeniculate connectivity, not changes in corticothalamic inputs, are responsible for the expression of SER-plasticity (Figures 2-3). However, we have previously shown that long-term disruption of cortical feedback activity can alter retinogeniculate connectivity during the experience-dependent phase of retinogeniculate development^9^. Therefore, it is possible that experience first alters corticothalamic activity, which subsequently triggers SER-plasticity by altering the weights of RGC inputs to the dLGN.

Plasticity in response to selective stimulus exposure has been demonstrated in the visual cortex of cats and mice^35,41–46^. While these changes are primarily attributed to local changes in cortex, our results highlight the potential contribution of earlier nodes of visual processing, as supported by recent work demonstrating ocular dominance plasticity in dLGN^30–34^. A recent study in ferrets has suggested the LGN does not contribute to rapid cortical plasticity following a 6 hour experience rearing protocol^47^—However, it is important to note that the rapid experience rearing protocol used to accelerate the maturation of direction-selectivity^48^ differs from that used in our study and in stripe rearing in cortex, which typically requires days to weeks to reveal an effect. Moreover, the mouse LGN has been likened to the primate and carnivore koniocellular/C LGN layers, which are underexplored and often not included in plasticity studies^1,49–51^. Future studies are needed to understand how plasticity differs across species and how the thalamus contributes to cortical function.

## Materials and Methods

### Animals

All procedures were approved by the Institutional Animal Care and Use Committee (IACUC) at Boston Children’s Hospital. Both male and female mice were used for all experiments. Mice were on a C57 BL/6J background (Jackson Laboratories strain #000664). Animals were reared from birth to P20 in normal rearing conditions with a 12:12 light-dark cycle. At P20, mice were weaned into separate cages and either placed in normal rearing conditions or in the dark room before undergoing selective experience rearing. For “recovery” experiments, SER mice were moved back into normal rearing conditions between P35-75. MeCP2 QKI mice^29^ were genotyped by standard PCR using the following primers. The mutant allele yields a 388bp product and the WT allele yields a 317bp product.

Forward: 5’– TCTAGCCCAATGACCCCCAA

Reverse: 5’– TCCCTGGGGACTGTAGAAACA

### Selective experience rearing (SER)

Animals were transported daily from the dark room to an optomotor chamber surrounded by 4 LED monitors. The home cage was placed inside the chamber and animals passively viewed the monitors for ∼3 hours per day before being transported back to the dark room. Between 2-4 mice were housed in a single cage during SER. Inside the chamber, mice were exposed to square-wave gratings moving at 2 cycles/second. The spatial frequency of the gratings was 0.04 cycles/degree measured from the center of the cage. The range of spatial frequencies animals experienced ranged from 0.025 - 0.057 cycles/degree depending on their location inside the cage. The mean luminance inside the chamber was 10 lux.

### In vivo electrophysiological recordings

Surgeries were performed while animals were under anesthesia (2% isoflurane in O_2_) and dexamethasone (2.5 mg/kg) was injected subcutaneously to prevent edema. Eyes were covered in silicone oil to prevent drying. A craniotomy that was 2-3mm in diameter was performed. For dLGN recordings, the craniotomy was centered ∼1.5mm anterior and ∼2.5mm lateral of the lambda suture. For visual cortex recordings, the craniotomy was centered ∼2.5mm lateral of the lambda suture. The exposed brain was always covered in artificial cerebral spinal fluid (aCSF) to prevent drying.

Prior to recording, chlorproxithene (0.3-0.5 mg/kg) was injected subcutaneously. Chlorproxithene is a sedative that permits isoflurane levels to be maintained at 0.5-1% during recordings. Lowered isoflurane levels result in more robust visual responses^23^. Previous studies have shown that anesthesia does not substantially alter orientation selectivity in the mouse dLGN^19,21^, therefore we opted to perform recordings in anesthetized animals for recording stability. The range of stereotaxic coordinates used for targeting dLGN were 0.8 - 1.6mm anterior and 2.10 - 2.35mm lateral from the lambda suture. Up to 4 penetrations were made per animal and penetrations were spaced at least 0.15mm apart in cartesian coordinates. The coordinates used for targeting visual cortex were 0.0 – 0.2mm anterior and 2.7 – 2.8mm lateral from the lambda suture. Only 1 penetration was made per animal for cortical recordings.

Recordings were made using linear 32-channel silicone probes (a1×32–25-5 mm-177, NeuroNexus and ASSY-37-H7b, Cambridge Neurotech). Data were acquired using a PZ5 digitizer with RZ5P processor (Tucker-Davis Technologies) with Synapse software running on a Workstation 8 computer (Tucker-Davis Technologies). Signals were sampled at 24414 Hz and filtered between 300 Hz and 5 kHz to isolate spikes for single-unit analysis. Recordings were always made from the right dLGN and the right eye was covered during visual stimulation. The electrode was initially advanced at a rate of ∼5 µm/second up to a depth of 2500µm. After reaching a depth of 2500µm, the animal was exposed to full-field drifting sine-wave gratings (0.04 cycles/degree, 2 cycles/second) and the electrode was advanced slowly (<1 µm/second). The electrode was lowered until the entire dorsal-ventral limits of dLGN were included in the 32-channel array. Upon reaching the final depth, the skull was covered with 2.5% agarose dissolved in aCSF and allowed to settle for at least 20 minutes prior to data collection.

To activate Gq-DREADDs in cortical silencing experiments, 3 mg/kg clozapine N-oxide (CNO) (Tocris) was injected subcutaneously. Because silencing occurred within 10 minutes (Figure 2C), animals were dosed with CNO at least 10 minutes prior to dLGN recordings. If recordings lasted longer than 2 hours, animals were re-dosed once. Recordings never lasted for more than 4 hours.

### Visual Stimuli

Visual stimuli were presented on a 21×12 inch LED monitor with a refresh rate of 60 Hz. The screen was positioned 22 cm away from the animal’s left eye offset at a 45-degree angle relative to the anteroposterior axis of the mouse. The screen covered ∼100° x 70° of visual space and the mean luminance of the screen was 20 lux. Stimuli were gamma corrected and generated using Psychophysics toolbox^52^ in MATLAB (MathWorks). To confirm that electrodes were in dLGN, contrast modulated white noise movies^23,24^ were presented to confirm that multi-unit receptive fields elevated as a function of ventral-dorsal axis along the linear electrode array^23^.

To determine the axis preference of dLGN neurons, full-field drifting sine-wave gratings consisting of 6 different spatial frequencies (0.01, 0.02, 0.04, 0.08, 0.16, and 0.32 cycles/degree) moving in 8 different directions were presented. Gratings were presented for 1.5 seconds with a temporal frequency of 2 cycles/second. Gratings conditions were presented in pseudorandom order over 12 trials and a blank gray screen condition was included to measure spontaneous activity. For experiments validating cortical silencing, full-field gratings with a spatial frequency of 0.04 cycles/degree and a temporal frequency of 2 cycles/second were presented. In a subset of experiments, full-field flashes were presented from darkness to determine the ON-OFF response polarity of dLGN units. Flashes consisted of 1s of light and dark alternating for 20 trials.

### Spike sorting and unit selection

Spike sorting was performed using default parameters in Kilosort2^53^. Kilosort outputs were manually curated in Phy (https://github.com/kwikteam/phy). To make curation as consistent and reproducible as possible, only clusters with a contamination percentage^54^ below 10% were selected: These clusters were automatically deemed “good” by Kilosort. Clusters labeled as multi-unit activity (mua) by Kilosort were excluded and clusters with a similarity index over 0.9 were automatically merged. Single units were manually excluded when they did not exhibit characteristic waveform shapes resembling extracellular action potentials or if the waveform amplitudes were not stable throughout the duration of the recording^54^. Additionally, units that were outside of the range of channels exhibiting clear multi-unit receptive fields used to map dLGN (see below) were excluded.

### Data analysis for responses to white noise movies

To confirm that penetrations were in dLGN, multi-unit spatial receptive fields were mapped during the experiment. Spike triggered averaging (STA) was performed on frames of the contrast modulated white noise movie presented 50 ms prior to each spike. For multi-unit receptive field mapping, spikes were identified as events that deviated 4 SDs below baseline. Only recordings from channels with clear multi-unit receptive fields that elevated along the ventral-dorsal axis of the electrode tract were included in subsequent analyses. For analysis of single-unit receptive fields, spike sorting was performed using Kilosort2 as described above. The spatial receptive fields of single units were fit with a 2D gaussian and only fits with a normalized mean square error < 0.1 were included for further analysis.

### Data analysis for responses to drifting gratings and flashes

Responses to gratings were calculated by measuring the mean firing rate (F0) in response to all unique spatial frequency and direction combinations. To determine if units were visual, responses to blank gray screen trials were compared with responses from the spatial frequency/direction combination that evoked the largest response. A Mann-Whitney test was performed on responses to blank trials vs. preferred trials and units were deemed visual if p < 0.01. The preferred spatial frequency was calculated by first measuring the direction that elicited the largest response. The spatial frequency that elicited the largest responses at the preferred direction was used as the preferred spatial frequency for each unit^19^.

To determine response reliability, a quality index (Qi) was calculated as previously described^55^. Spiking responses were binned into 0.1s bins to create a Trial x Response matrix, which represents gratings responses to each unit’s optimal frequency/direction over multiple trials. Qi was calculated by dividing the variance of the mean response by the mean variance across trials. Units with a Qi < 0.25 were excluded (Figure S3). Axis-selectivity index (ASI) was computed as (R_preferred_ – R_orthogonal_) / (R_preferred_ + R_orthogonal_), where R_preferred_ is the mean response to gratings moving along the preferred axis. R_orthogonal_ is the mean response to gratings moving along the axis orthogonal to R_preferred_. Units with an ASI > 0.33 were classified as AS. Analyses were also performed measuring circular variance to calculate “global” ASI (gASI); These analyses yielded identical conclusions.

The preferred axes of AS units were calculated from the vector sum of average responses to each direction in orientation space and the angle of the resultant vector was computed^56^. To determine whether the distribution of preferred axes significantly differed between rearing conditions, the preferred axes of AS units was divided into 3 groups (horizontal, vertical and diagonal). These groups were chosen because animals were presented with gratings moving in 8 directions (0-315 degrees evenly spaced in 45 degree increments). Thus, units could be categorized as horizontal, vertical, or diagonal preferring based on the gratings that elicited maximal response. Alternatively, units could be categorized by binning preferred axes computed by vector sum into 3 groups. These two methods yielded in identical results. A chi-squared (Χ^2^) test was then used to determine whether SER altered the distribution of preferred axes. P-values < 0.05 were considered significant.

ON-OFF index of units in response to full-field flashes was computed as follows: (R_ON_ - R_OFF_) / (R_ON_ + R_OFF_). R_ON_ and R_OFF_ represent the mean firing rate of each unit calculated during the first 400ms following light onset (R_ON_) and offset (R_OFF_). All analyses were performed without knowledge of the rearing condition. During recordings, experimenters were blind to the rearing condition of the mouse in most cases (55/91 animals).

### Neonatal stereotaxic injections

For acute cortical suppression with DREADDs, viral injections into cortex were performed in mouse pups between P1-3 to ensure sufficient viral expression by P27-35. Excitatory (Gq) DREADDs were expressed in inhibitory neurons by injecting AAV9-hDlx-GqDREADD-P2A-dTomato (Addgene #83897-AAV9, 2.1 x 10^13^ vg/mL) unilaterally. Subsequent dLGN recordings were performed on the same side as the cortical injection. Injections were made in C57 pups anesthetized by hypothermia and head fixed on a chilled stereotaxic adaptor for neonatal mice (Stoelting, Catalog #: 51625). Virus was loaded in glass micropipettes and injected using a Nanoject III microinjector (Drummond Scientific) along 3 sites along the medial-lateral axis.

These sites were approximately 500, 900 and 1300µm lateral from the midline and 100-200µm anterior to the lambda suture. 50nL injections were made at 2 different depths (120 and 250µm) at a rate of 1nL/second. The dTomato signal was amplified using immunohistochemistry (described below) to confirm viral expression across the entire extent of visual cortex.

Expression in subcortical areas was never observed because the Dlx promoter restricts expression to cortex^28^. To cover as much of visual cortex as possible, injections typically spanned hippocampus and adjacent non-visual cortical areas.

### Histology

After *in vivo* recordings, the brain was dissected and drop fixed in 4% paraformaldehyde (Electron Microscopy Sciences) in 1X PBS at 4°C overnight. 100µm coronal sections were made using a Leica 1000S vibratome. Sections were incubated in blocking solution (6% goat serum and 0.1% Triton in 1X PBS) overnight at 4°C. Sections were then incubated for 1-2 days at 4°C in primary antibody solution containing rabbit anti-RFP (1:1000–1:2000, Rockland Catalog #: 600-401-379) and mouse anti-NeuN (1:500-1:1000, Millipore Catalog #MAB377) in 3% goat serum and 0.1% Triton in 1X PBS. Then, sections were washed in 1X PBS and incubated overnight at 4°C in secondary antibody solution containing Alexa 555 goat anti-rabbit (1:1000, Thermo Fisher Catalog #: A-21428). After washing in 1X PBS, sections were mounted on glass slides and coverslipped using Fluoromount Aqueous Mounting Medium (Millipore Sigma).

To anatomically confirm electrode penetrations and viral expression in cortical silencing experiments, slides were imaged using an epifluorescence microscope (Zeiss Axio Imager). Electrode placement within the dLGN was confirmed using multi-unit receptive field mapping as described above. AS units were found across the entire depth of dLGN (data not shown). In a subset of experiments (19/72 animals), penetrations were also confirmed anatomically.

### Exclusion Criteria and Statistics

Recordings were excluded if 1) the multi-unit receptive fields recorded during a penetration did not follow predicted retinotopic organization^23^, 2) if no “good” single units were found by Kilosort (see above) and 3) if fewer than 4 responsive axis-selective units were recorded from a given animal. For cortical silencing experiments, animals were excluded if extensive viral labeling was not observed in visual cortex. 8 animals were excluded in this study due to poor viral expression of Dlx-GqDREADD virus. Power analyses to estimate sample sizes for Χ^2^ tests were performed using G*Power^57^. Specific information about statistical tests is included in the figure legends and the main text. p < 0.05 was considered significant.

---

**Figure S1:**
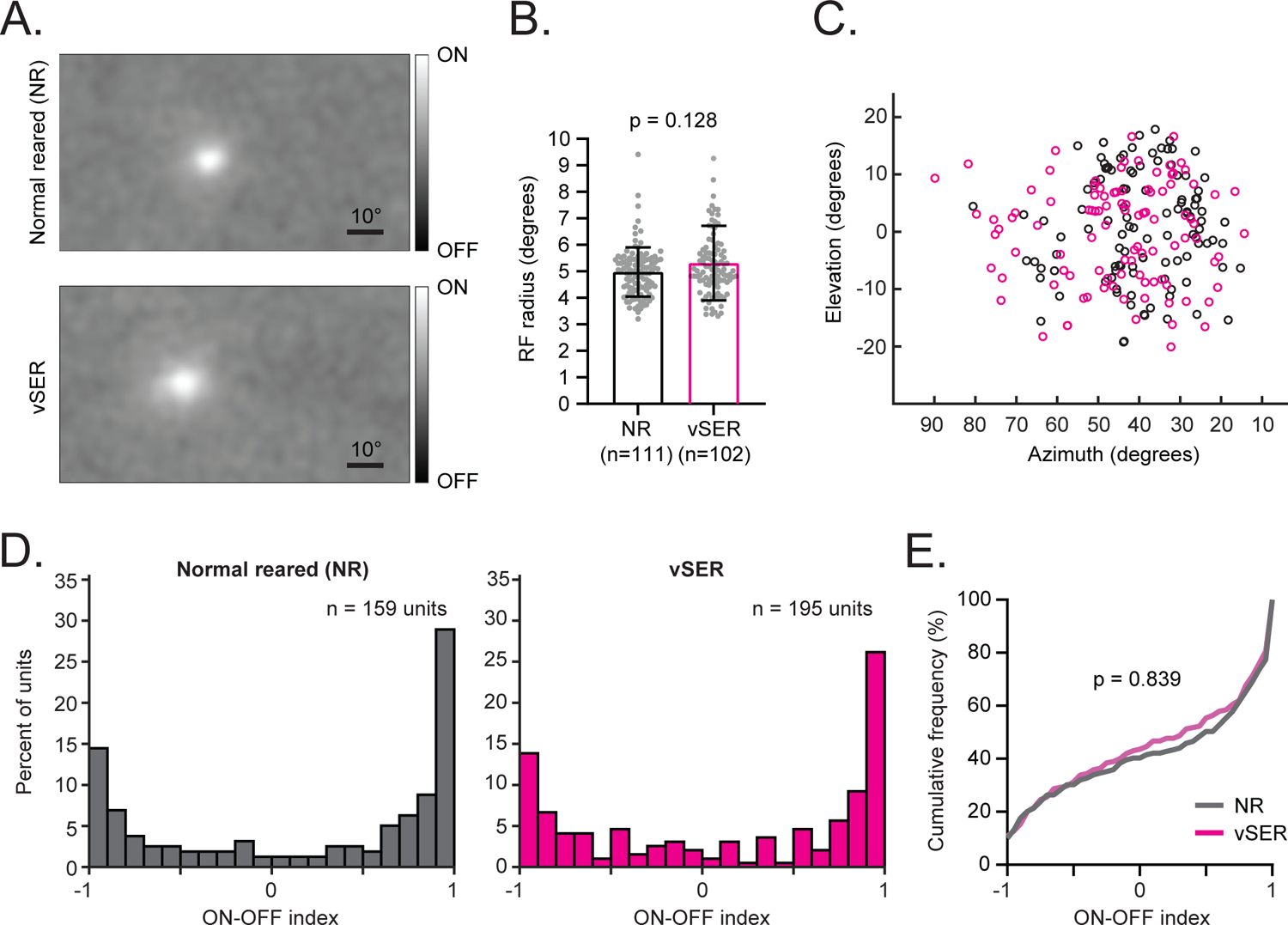
dLGN receptive field sizes and ON-OFF responses are not significantly altered by SER. **(A)** Examples of receptive fields (RFs) measured from dLGN single units in normal reared (top) and vSER (bottom) animals. RFs were acquired by spike-triggered averaging contrast modulated noise movies with a 50ms time lag. **(B)** Grouped data showing RF sizes of single units in the dLGN between P27-35 in normal reared (black) and vSER (magenta) animals. There were no significant differences in dLGN RF sizes in NR and vSER animals (Mann-Whitney test, p = 0.128). **(C)** RF locations of single units in dLGN recorded in NR (gray) and vSER (magenta) animals. **(D)** Histograms showing the distribution of ON-OFF index in units that were visually responsive to full-field light flashes. dLGN recordings are from a subset of NR (gray, n = 4 animals) and vSER (magenta, n = 6 animals) recordings used in Figure 1 because full-field flashes were not always included in the visual stimulation protocol. **(E)** Same data as panel (D) but plotted as cumulative distribution. There was no significant change in the distribution of ON-OFF index in response to vSER (Kolmogorov-Smirnov test, NR vs. vSER p = 0.839).

**Figure S2:**
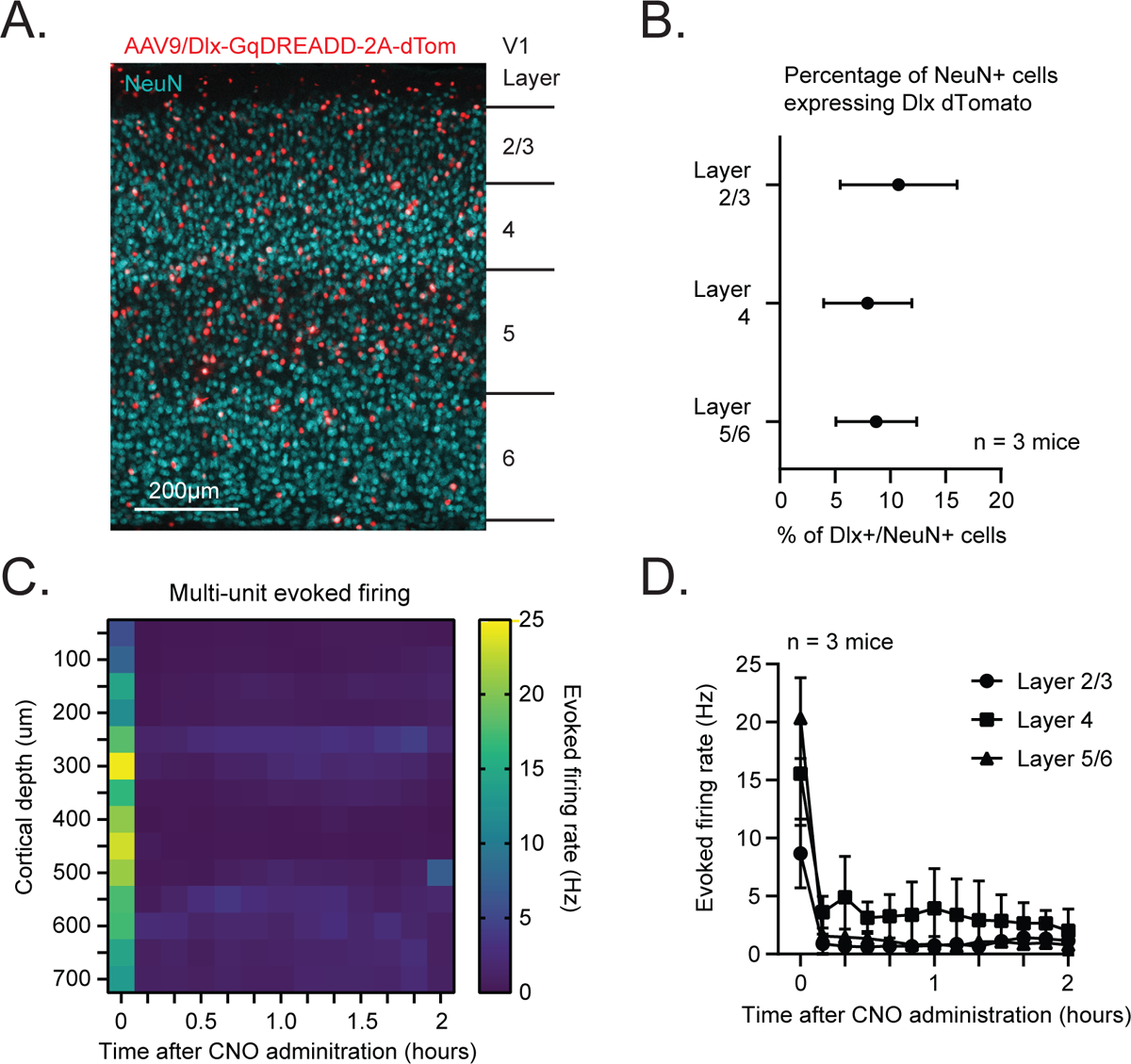
Silencing across cortical depths with Dlx-GqDREADD virus. **(A)** Example coronal section of the visual cortex from an animal that received cortical injection of AAV9/Dlx-GqDREADD-2A-dTomato between P0-3 and sacrificed at P35. The section was immunolabeled for NeuN (cyan) to visualize neurons and red fluorescent protein to visualize cells expressing dTomato (red). **(B)** Grouped data showing the percentage of neurons (NeuN+) that were dTomato+ across different layers of visual cortex. **(C)** Heat maps from an example recording showing evoked firing across multielectrode array that spanned 700um of visual cortex. Each square represents the average multi-unit firing rate from a single recording site over 10 minutes. Evoked firing was measured as responses to full-field drifting gratings. Time 0 represents baseline measurements taken prior to systemic CNO administration. **(D)** Grouped data showing mean evoked firing rate in different cortical layers after systemic CNO administration. Time 0 represents baseline measurements taken prior to CNO administration.

**Figure S3:**
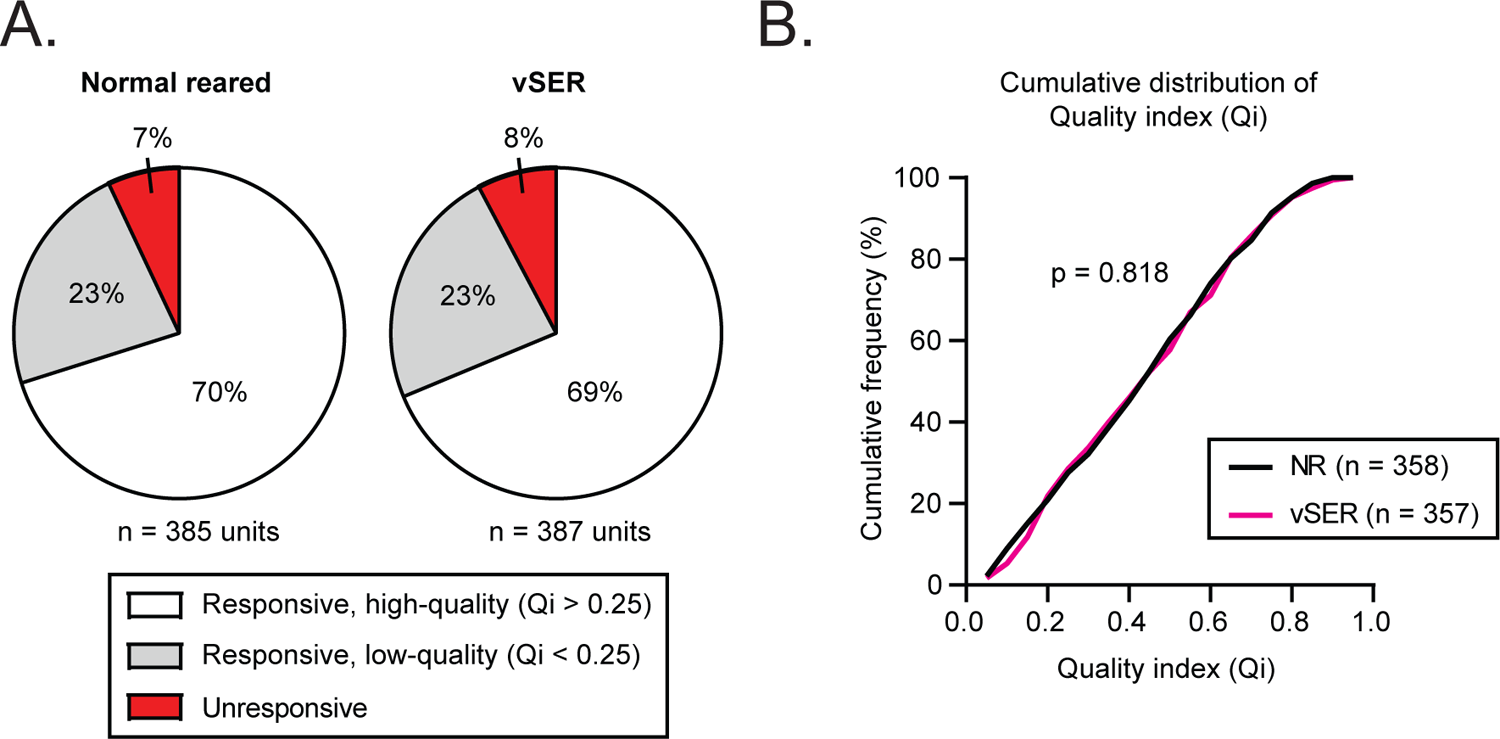
The proportion of visually responsive units is not altered by SER. **(A)** Proportion of units that were visually unresponsive (red), visually responsive with low-quality responses (gray) and visually responsive with high-quality responses (white). SER did not significantly alter the proportion of visually responsive units (Χ^2^ test, p = 0.837). Units are from the same dataset as Figure 1E-H. **(B)** Cumulative distribution of Qi in dLGN units recorded in normal reared (NR, black) and vSER (magenta) animals. vSER did not significantly alter the quality of dLGN recordings (Kolmogorov-Smirnov test, p = 0.818).

